# Yap regulates glucose utilization and sustains nucleotide synthesis to enable organ growth

**DOI:** 10.1101/300053

**Authors:** Andrew G. Cox, Allison Tsomides, Dean Yimlamai, Katie L. Hwang, Joel Miesfeld, Giorgio G. Galli, Brendan H. Fowl, Michael Fort, Kimberly Y. Ma, Mark R. Sullivan, Aaron M. Hosios, Erin Snay, Min Yuan, Kristin K. Brown, Evan C. Lien, Sagar Chhangawala, Matthew L. Steinhauser, John M. Asara, Yariv Houvras, Brian Link, Matthew G. Vander Heiden, Fernando D. Camargo, Wolfram Goessling

**Author notes:** Present-address: Organogenesis and Cancer Program, Peter MacCallum Cancer Centre, Victoria, Australia; Sir Peter MacCallum Department of Oncology and Department of Biochemistry and Molecular Biology, The University of Melbourne, Victoria, Australia (Andrew G. Cox). Present-address: Department of Pediatrics, Children’s Hospital of Pittsburgh, University of Pittsburgh Medical Center, Pittsburgh, PA. Present-address: Novartis Institutes for BioMedical Research, Disease Area Oncology, 4056 Basel, Switzerland (Giorgio G. Galli). Present-address: Cancer Therapeutics Program, Peter MacCallum Cancer Centre, Victoria, Australia; Sir Peter MacCallum Department of Oncology and Department of Biochemistry and Molecular Biology, The University of Melbourne, Victoria, Australia (Kristin K. Brown). Correspondence (AGC) (WG).

## Abstract

The Hippo pathway and its nuclear effector Yap regulate organ size and cancer formation. While many modulators of Hippo activity have been identified, little is known about the Yap target genes that mediate these growth effects. Here, we show that *yap*-/- mutant zebrafish exhibit defects in hepatic progenitor potential and liver growth due to impaired glucose transport and nucleotide biosynthesis: transcriptomic and metabolomic analyses reveal that Yap regulates expression of glucose transporter *glut1*, causing decreased glucose uptake and use for nucleotide biosynthesis in *yap*-/- mutants, and impaired glucose tolerance in adults. Nucleotide supplementation improved Yap-deficiency phenotypes, indicating functional importance of glucose-fuelled nucleotide biosynthesis. Yap-regulated *Glut1* expression and glucose uptake are conserved in mammals, suggesting that stimulation of anabolic glucose metabolism is an evolutionarily conserved mechanism by which the Hippo pathway controls organ growth. Together, our results reveal a central role for Hippo signalling in metabolic homeostasis.

## Introduction

Control of size at the cell, organ, and organismal level is of fundamental biological importance. The Hippo pathway has recently emerged as a master regulator of organ size control. Hippo pathway signalling is modulated by the integration of environmental cues, including cell-cell contact, cell polarity, G protein-coupled receptor stimulation, mechanical forces and nutrient status (Harvey et al. 2013; Piccolo et al. 2014; Yu et al. 2015). These stimuli impact the activity of the core Hippo kinase cascade, thereby regulating nuclear localization of the transcriptional co-activators Yap and Taz, which bind to the Tead family of transcription factors to stimulate tissue growth (Harvey et al. 2013; Piccolo et al. 2014). Prior murine studies have demonstrated that conditional expression of activated Yap in the liver stimulates hepatomegaly and drives tumorigenesis (Camargo et al. 2007; Dong et al. 2007; Zhou et al. 2009; Lu et al. 2010; Yimlamai et al. 2014). It is now recognized that many oncogenic pathways modulate the Hippo pathway and sustain tumor growth in a Yap-dependent manner (Harvey et al. 2013; Piccolo et al. 2014; Yu et al. 2015). Much less is known, however, about the requirement for Yap for normal organ development and tissue growth, as global knockout murine models show early embryonic lethality (Morin-Kensicki et al. 2006). Further, the transcriptional program regulated by Yap to control organ growth is incompletely understood.

Altered cellular metabolism is a hallmark of cancer cells (Hanahan and Weinberg 2011). Tumor cells exploit metabolic pathways to enhance nutrient uptake and fulfill the biosynthetic requirements needed to increase biomass (Howell et al. 2013; Mayers and Vander Heiden 2015). Aerobic glycolysis (“Warburg effect”) is the best described metabolic change in cancer (Vander Heiden et al. 2009). Recent studies have identified additional mechanisms by which oncogenic pathways reprogram metabolism in cultured cancer cells (Mayers and Vander Heiden 2015). In addition, we have recently demonstrated that the Hippo pathway effector Yap reprograms glutamine metabolism *in vivo* to stimulate nucleotide biosynthesis and promote liver growth during development and tumorigenesis (Cox et al. 2016a). Given the profound impact of Yap on nitrogen metabolism, we hypothesize that Yap enables the cell to change other aspects of its energy metabolism to provide the building blocks for sustained growth during development and organ homeostasis.

In this study, we perform a combination of transcriptomic and metabolomic analyses in *yap* mutant and transgenic zebrafish to discover that Yap directly stimulates cellular glucose uptake and anabolic glucose utilization for *de novo* nucleotide biosynthesis through induction of the glucose transporter Glut1. *yap*-/- mutant zebrafish exhibit defects in liver progenitor formation and organ growth during development, and glucose intolerance in adulthood; in contrast, activation of Yap increases liver size. Exposure to GLUT inhibitors suppresses Yap-induced liver growth, whereas nucleotide supplementation alleviates defects in *yap*-/- mutants. Importantly, regulation of *Glut1* and glucose uptake by Yap is conserved in mammals. Together, our findings define Yap as a novel regulator of anabolic glucose metabolism that supports the nucleotide synthesis requirements for organ growth.

## Results

### Yap is required for optimal liver development

The impact of Yap on organogenesis has been difficult to address in mammalian models due to early embryonic lethality of global Yap knockout mice (Morin-Kensicki et al. 2006). To investigate the role of Yap in liver development, we examined *yap*-/- mutant zebrafish embryos (**Fig.S1A**) (Miesfeld et al. 2015), which exhibited multiple developmental abnormalities, ranging from eye pigmentation defects in mildly affected embryos to cardiac edema in more severely affected embryos. In contrast to knockout mice, ~50% of *yap*-/- mutants survived into adulthood (**Fig.S1B,C**). Liver growth in *yap*-/- mutants, examined in a fluorescent hepatocyte reporter line (*Tg(*-*2.8fabp10a:CFP*-*NTR)*, hereinafter referred to as *lf:CFP*) (Choi et al. 2014), was impaired, revealing a ~32% reduction in fluorescent liver area at 3 and 5 days post fertilization (dpf) compared to wild-type (WT) siblings (**Fig.1A,B; S1D,E**). Whole mount *in situ* hybridization (WISH) analysis confirmed impaired liver growth and demonstrated that growth of other endodermally-derived tissues, such as intestine (*intestinal fatty acid binding protein*, *ifabp*) and exocrine pancreas (*trypsin*), was also impaired by loss of *yap* (**Fig.S1F**). The livers of *yap*-/- mutant larvae exhibited abnormal histological features (hepatocyte ballooning and sinusoidal widening) with a reduced number of hepatocytes compared to WT (**Fig.1A; S1G**). Together, these studies demonstrate that *yap* deficiency impairs liver growth during development.

**Figure 1:**
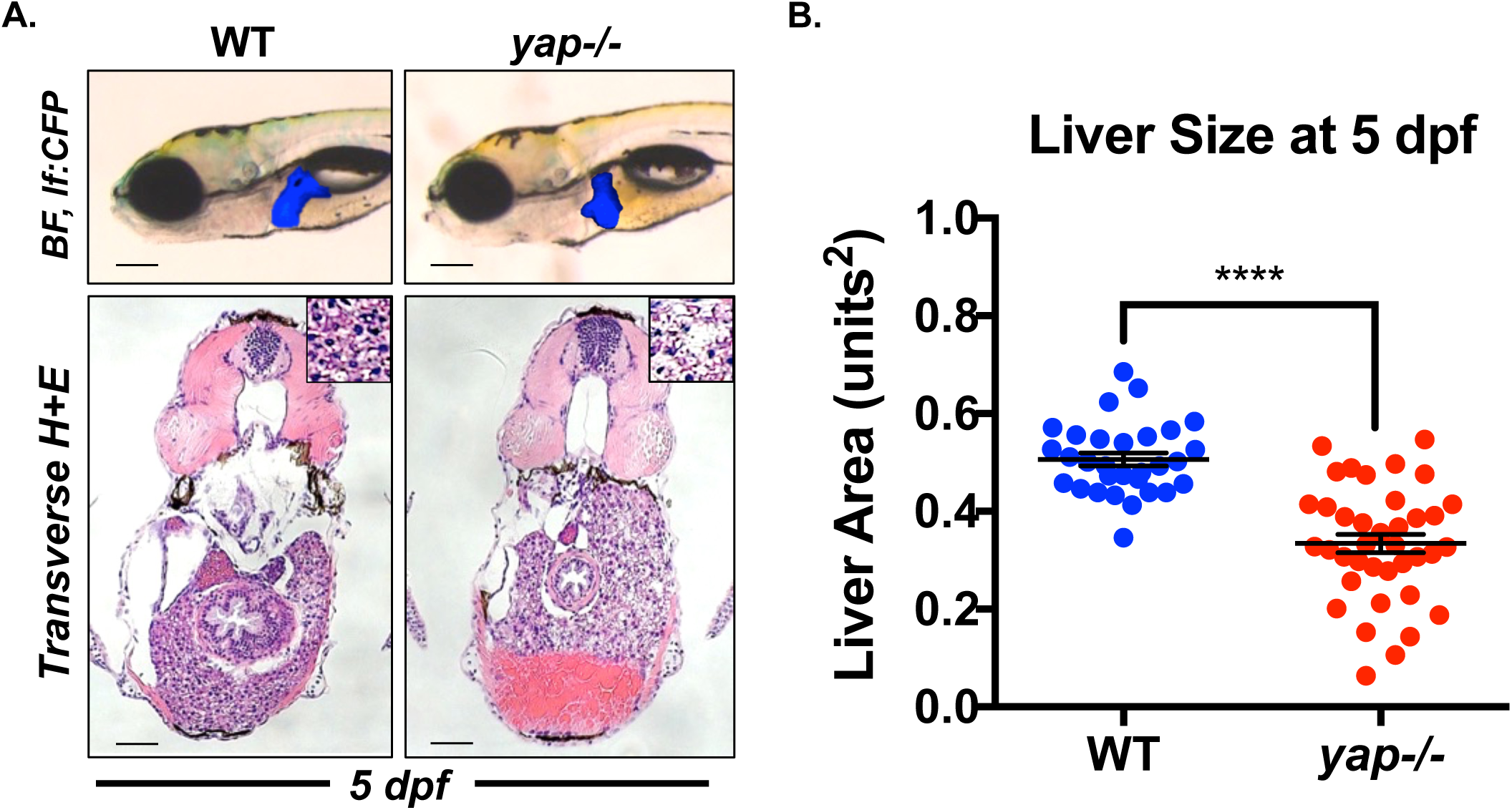
The Hippo effector Yap is required for optimal liver development. A) Fluorescent and histological analysis of liver morphology in WT and *yap*-/- mutant larvae on a *lf:CFP* background at 5 dpf. The blue colour overlay represents the fluorescent liver tissue. Scale bar: 200μm (bright field, BF) and 50μm (histology, H+E). B) Quantitative analysis of fluorescent liver area in WT and *lf:Yap* larvae at 5 dpf. n>30, ^∗∗∗∗^p<0.0001, two-sided Student’s t-test; values represent the mean ± standard error of the mean (s.e.m.).

### Yap regulates hepatoblast formation and expansion

To assess whether impaired liver development in *yap*-deficient embryos could be caused by defects in hepatoblast formation, WISH and qPCR analyses were performed, revealing significantly reduced expression of hepatoblast markers *prox1* and *hhex* in *yap*-/- embryos at 36 hours post fertilization (hpf) (**Fig.2A,B**). To further elucidate the developmental impact of Yap, we generated transgenic zebrafish conditionally expressing activated Yap (Tg*(hsp70:mcherry*-*2A*-*flag*-*yap1^S87A^)*, referred to as *hs:Yap*) or a dominant-negative form of Yap (Tg*(hsp70:dnyap*-*2A*-*mcherry)*, referred to as *hs:dnYap)* under a heat-shock promoter (**Fig.S2A**), enabling the precise temporal regulation of Yap activity. Detection of *mcherry*, *ctgf* and *amotl2b* by qPCR confirmed effective induction or suppression of Yap target genes upon heat schock (**Fig.S2B,C,D**). Induction of dominant-negative Yap at 24 hpf during liver specification suppressed *prox1* and *hhex* expression at 36 hpf, whereas induction of activated Yap increased *prox1* and *hhex* expression areas (**Fig.2C,D**). Collectively, these data demonstrate that Yap regulates liver progenitor development.

**Figure 2:**
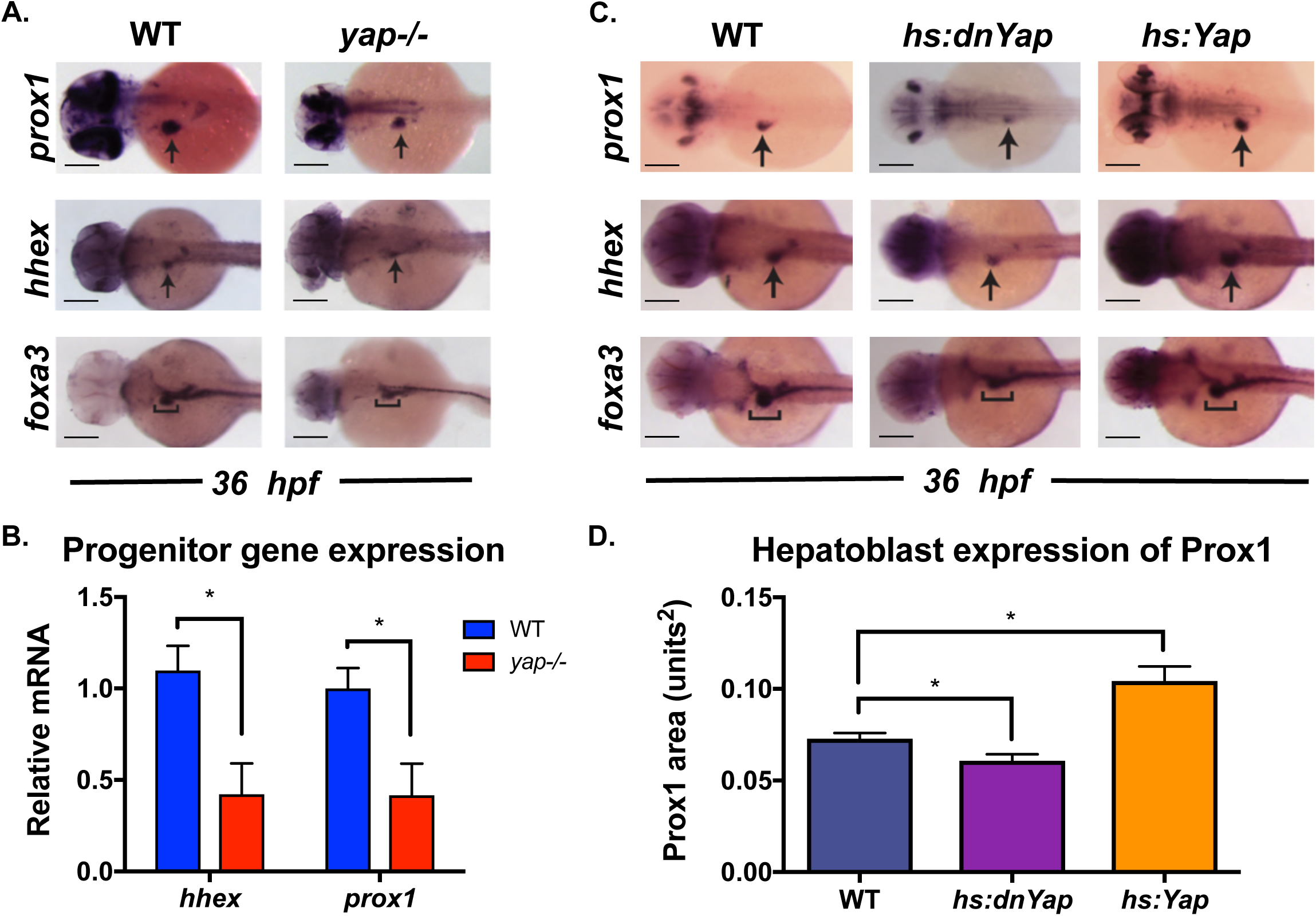
Yap regulates hepatoblast formation and expansion. A) Whole-mount in situ hybridization (WISH) analysis of progenitor marker *prox1*, *hhex*, and *foxa3* expression in WT and *yap*-/- mutant embryos at 36 hpf. Arrow and bracket highlight the hepatic bud. Scale bar: 200μm. B) qPCR analysis of *hhex* and *prox1* gene expression in WT and *yap*-/- mutant embryos at 48 hpf. n>3, ^∗^p<0.05, two-sided Student’s t-test; values represent the mean ± s.e.m. C) WISH analysis of *prox1*, *hhex* and *foxa3* expression in WT, *hs:dnYap* and *hs.Yap* transgenic embryos heat shocked at 24 hpf and fixed at 36 hpf. Arrow and bracket highlight the hepatic bud. Scale bar: 200μm. D) Quantitative analysis of hepatic bud area (*prox1* staining) in WT, *hs:dnYap* and *hs.Yap* transgenic embryos heat shocked at 24 hpf and fixed at 36 hpf. n>10, ^∗^p<0.05, two-sided Student’s t-test; values represent the mean ± s.e.m.

### Yap deficient adult zebrafish exhibit liver hypoplasia

Adult *yap*-/- mutants were easily identifiable from WT fish due to their smaller size (**Fig.S3A**), while dissection revealed smaller livers compared to WT fish (**Fig.3A**). Histological examination demonstrated decreased hepatocyte size of *yap*-/- hepatocytes (**Fig.3A**). *yap*-/- mutants had a significantly lower body mass, and further exhibited a 15% reduction in their liver:body mass ratios compared to WT, indicating a specific effect on liver growth independent of, or in addition to, decreased overall body size (**Fig.3B,C**). *yap*-/- mutant livers also contained reduced glycogen content, as demonstrated by PAS staining, without signs of fibrosis (Masson’s Trichrome stain) or endothelial dysfunction (Reticulin stain) (**Fig.S3B**). Given that Yap regulated hepatoblasts during development, we examined how Yap deficiency affected hepatic progenitors in adulthood. Expression levels of hepatoblast markers *hhex* and *prox1* were reduced by 80% and 55%, respectively, in adult *yap*-/- mutant liver tissue (**Fig.3D,E**). These studies reveal that *yap* deficiency in adulthood leads to liver hypoplasia and a reduced pool of hepatic progenitors.

**Figure 3:**
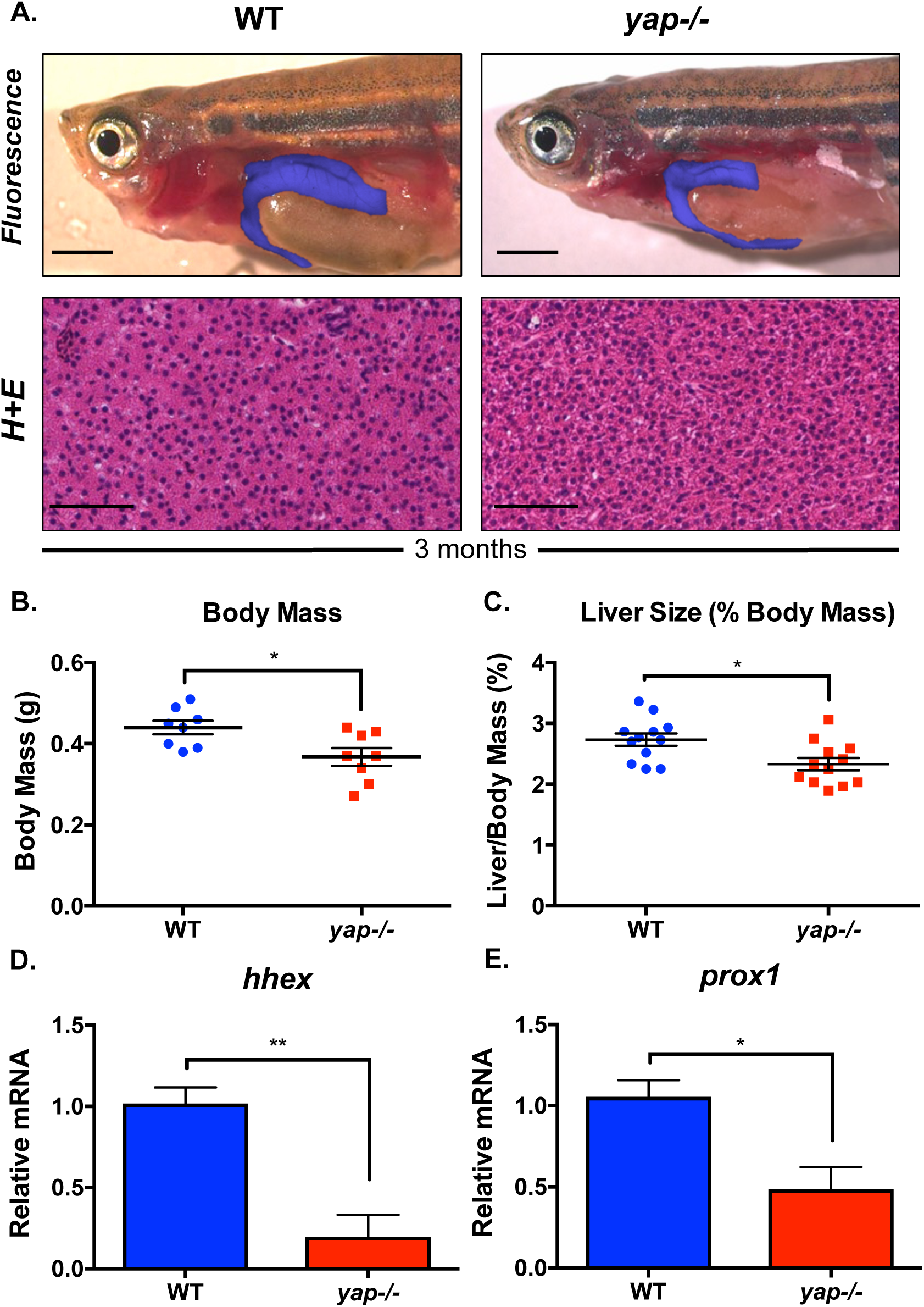
Yap deficient adults are small and exhibit liver hypoplasia. A) Fluorescent and histological analysis of liver morphology in dissected WT and *yap*-/- mutant adults on a *lf:CFP* background. The blue colour overlay represents the fluorescent liver tissue. Scale bar: 2mm (brightfield, BF) and 50μm (histology, H+E). B) Quantitative determination of body mass in WT and *yap*-/- adults. n=8, ^∗^p<0.05, two-sided Student’s t-test; values represent the mean ± s.e.m. C) Quantitative determination of liver size in WT and *yap*-/- adults as determined by measuring liver mass as a percentage of body mass. n=12, ^∗^p<0.05, two-sided Student’s t-test; values represent the mean ± s.e.m. D) qPCR analysis of the *hhex* expression in adult liver tissue isolated from WT and *yap*-/- fish. n>3, ^∗∗^p<0.01, two-sided Student’s t-test; values represent the mean ± s.e.m. E) qPCR analysis of the *prox1* expression in adult liver tissue isolated from WT and *yap*-/- mutant fish. n>3, ^∗^p<0.05, two-sided Student’s t-test; values represent the mean ± s.e.m.

### Yap regulates expression of *glut1* and *glut2*

To identify Yap target genes responsible for the observed defects in liver growth, transcriptomic analysis was accomplished by RNAseq in WT and *yap*-/- mutant embryos at 3 dpf, revealing a large subset of differentially expressed genes (**Fig.4A**). Cross-examination of these datasets was performed with differentially expressed genes in *yap*-/-*;taz*-/- double mutant embryos, as well as previously analyzed adult *lf:Yap* liver tissue (Cox et al. 2016a), revealing 191 genes that, compared to WT, were significantly downregulated in both *yap*-/- mutant and *yap*-/-*;taz*-/- mutants and induced in *lf:Yap* embryos (**Fig.S4A**). Among these 191 genes, the glucose transporters *glut1* and *glut2* were significantly downregulated in *yap*-/- mutants (**Fig.4B; S4B**). qPCR analysis of *yap*-/- embryos at 3 dpf confirmed reduction in *glut1* and *glut2* expression by 84% and 71%, respectively (**Fig.4C**). *glut1* and *glut2* expression in the heat-shock inducible transgenic lines was significantly reduced upon induction of *hs:dnYap* at 48 hpf, whereas *hs:Yap* embryos expressed elevated levels of *glut1* and *glut2* (**Fig.4D**). Finally, glucose transporter expression was suppressed in adult *yap*-/- mutant liver tissue, while *lf:Yap* transgenic livers contained 2-3 fold elevated levels of *glut1* and *glut2* (**Fig.4E**). Together, these experiments demonstrate that Yap is necessary and sufficient to regulate expression of *glut1* and *glut2.*

**Figure 4:**
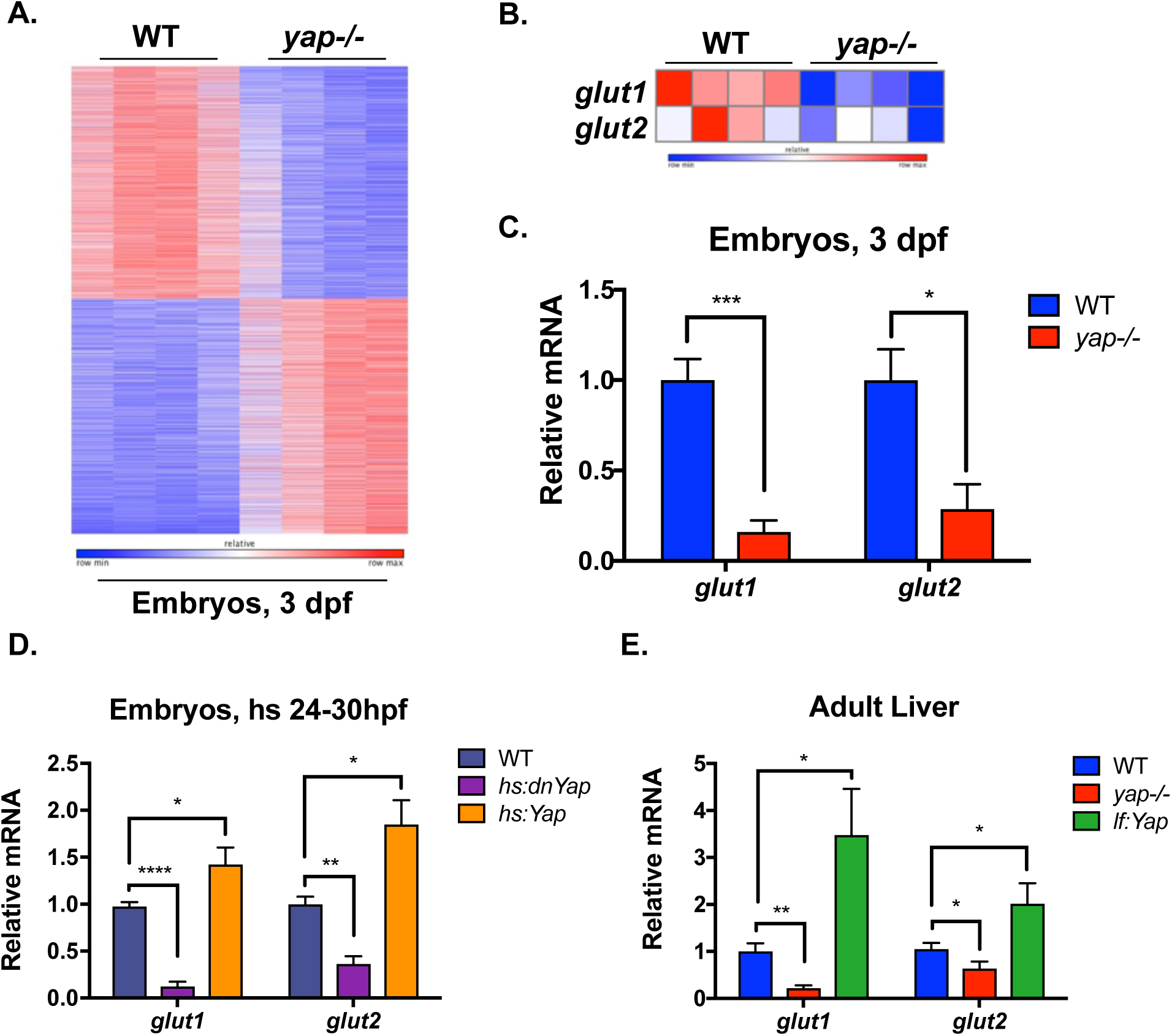
Transcriptional profiling reveals that Yap regulates the expression of *glut1* and *glut2*. A) Heatmap of RNAseq analysis illustrating statistically significant differential gene expression between WT and *yap*-/- mutant larvae at 3 dpf. Blue indicates decreased expression, while red demonstrates increased expression. B) Heatmap analysis of RNAseq data highlights the reduction of glucose transporters (*glut1* and *glut2*) in *yap*-/- mutant larvae at 3 dpf. C) qPCR analysis of *glut1* and *glut2* expression in WT and *yap*-/- mutant larvae at 3 dpf. n>3, ^∗^p<0.05, ^∗∗∗^p<0.001. D) qPCR analysis of *glut1* and *glut2* expression in WT, *hs:dnYap* and *hs:Yap* transgenic embryos heat shocked at 24 hpf and isolated at 30 hpf. n>3, ^∗^p<0.05, ^∗∗^p<0.01, ^∗∗∗∗^p<0.0001, two-sided Student’s t-test; values represent the mean ± s.e.m. E) qPCR analysis of *glut1* and *glut2* expression in liver tissue dissected from WT, *yap*-/- mutant and *lf:Yap* transgenic adults. n>3, ^∗^p<0.05, ^∗∗^p<0.01, two-sided Student’s t-test; values represent the mean ± s.e.m.

### Yap regulates glucose metabolism and nucleotide biosynthesis

Given that Yap regulates the expression of glucose transporters, we examined the impact of Yap loss on glucose homeostasis in adults, using a glucose tolerance test (GTT). In WT fish, serum glucose levels tripled within 60 mins after intraperitoneal injection of a weight-based glucose dose and normalized by 3-4 hours post injection. These findings indicate a previously unrecognized highly conserved regulation of glucose homeostasis compared to mammals. *yap*-/- mutants at 3 months of age exhibited glucose intolerance, with a ~25% reduced rate of glucose clearance from the blood into peripheral tissue at 120-240 min (**Fig.5A**). Surprisingly, glucose intolerance was markedly exacerbated in aged *yap*-/- mutant fish at 12 months, where glucose levels did not decrease over 4 hours (**Fig.5B**), indicating metabolic dysregulation in *yap*-/- mutant fish as they age. Together, these studies suggest that Yap plays an important role in glucose homeostasis.

**Figure 5:**
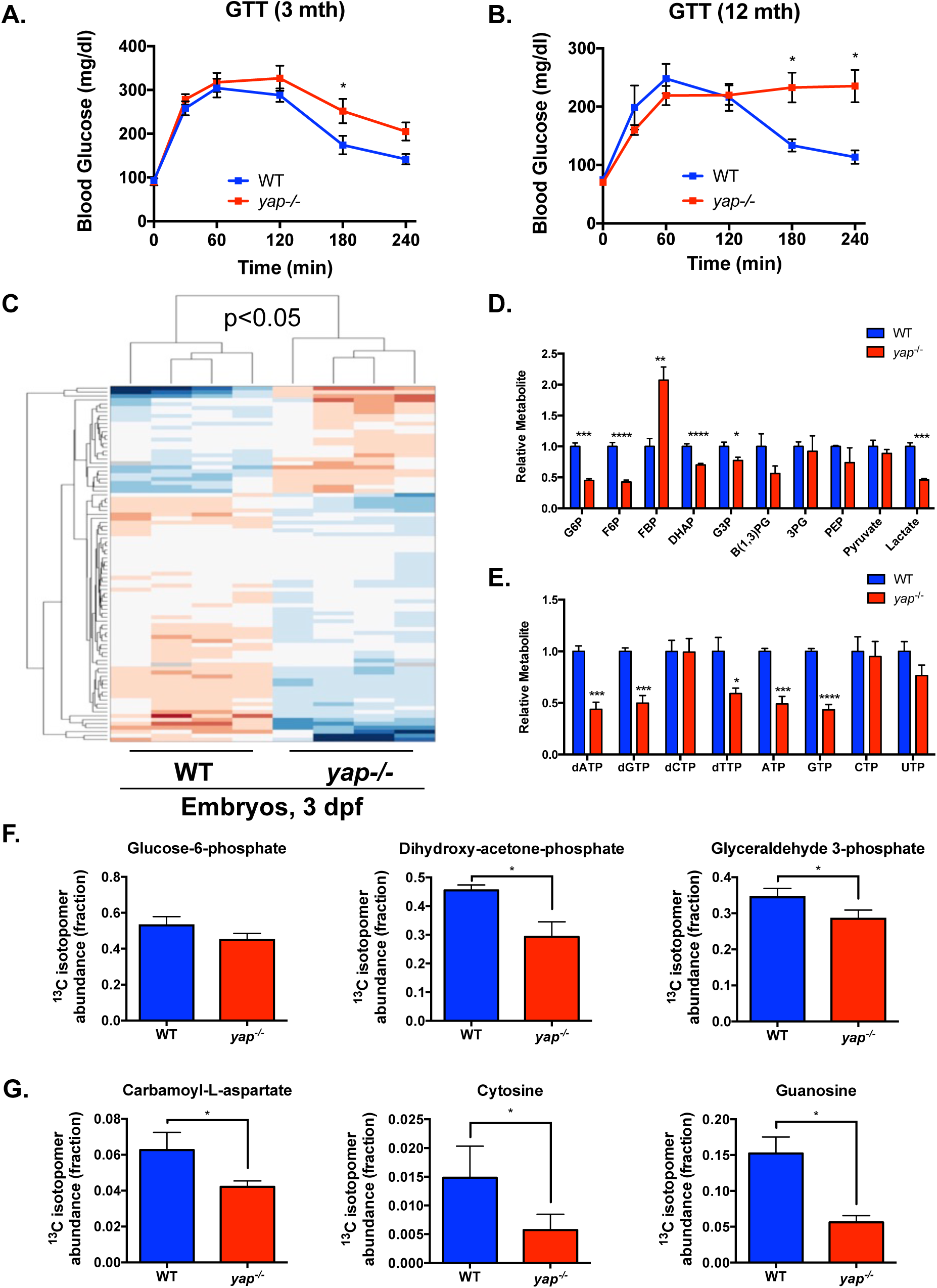
Yap regulates glucose metabolism and nucleotide biosynthesis. A) Glucose tolerance test in 3-month old WT and yap-/- mutant zebrafish. Blood glucose was measured 0, 30, 60, 120, 180 and 240 min post glucose injection. n>8, ^∗^p<0.05, two-sided Student’s t-test; values represent the mean ± s.e.m. B) Glucose tolerance test in 12-month old WT and yap-/- mutant zebrafish. Blood glucose was measured 0, 30, 60, 120, 180 and 240 min post glucose injection. n>8, ^∗^p<0.05, two-sided Student’s t-test; values represent the mean ± s.e.m. C) Clustergram analysis of polar metabolite abundance from WT and *yap*-/- mutant larvae at 3 dpf as determined by LC-MS/MS via selected reaction monitoring (SRM) analysis. n=4, p<0.05. D) Steady state abundance of glycolytic intermediates in WT and *yap*-/- mutant larvae as determined by SRM analysis. n=4, ^∗^p<0.05, ^∗∗^p<0.01, ^∗∗∗^p<0.001, ^∗∗∗∗^p<0.0001, two-sided Student’s t-test; values represent the mean ± s.e.m. E) Steady state abundance of nucleotides in WT and *yap*-/- mutant larvae as determined by SRM analysis. n=4, ^∗^p<0.05, ^∗∗∗^p<0.001, ^∗∗∗∗^p<0.0001, two-sided Student’s t-test; values represent the mean ± s.e.m. F) Relative isotopic enrichment of ^13^C in glycolytic intermediates following incubation of WT and *yap*-/- mutant larvae in [U^13^C_6_]-glucose as determined by SRM analysis. n=3, ^∗^p<0.05, two-sided Student’s t-test; values represent the mean ± s.e.m. G) Relative isotopic enrichment of ^13^C in nucleotide precursors following incubation of WT and *yap*-/- larvae in [U^13^C_6_]-glucose as determined by SRM analysis. n=3, ^∗^p<0.05, two-sided Student’s t-test; values represent the mean ± s.e.m.

To further illumniate glucose utilization in *yap*-/- mutants, polar metabolomics analysis with targeted liquid chromatography – tandem mass spectrometry (LC/MS-MS) via selected reaction monitoring (SRM) was performed (Yuan et al. 2012). Hierarchical clustering of metabolite abundance revealed a distinctive metabolic state in *yap*-/- mutant embryos with significant alterations in steady-state levels of metabolites enriched in glycolysis and *de novo* nucleotide biosynthesis (**Fig.5C; S5A**). Specifically, *yap*-/- embryos contained significantly reduced levels of glycolytic intermediates glucose-6-phosphate (G6P), fructose-6-phosphate (F6P), dihydroxyacetone phosphate (DHAP), glyceraldehyde-3-phosphate (G3P) and lactate, as well as purine nucleotides deoxyadenosine triphosphate (dATP), deoxyguanosine triphosphate (dGTP), adenosine triphosphate (ATP) and guanosine triphosphate (GTP) (**Fig.5D,E**). The decrease of nucleotide triphosphates in *yap*-/- mutants coincided with an increase in adenosine monophosphate (AMP), guanosine monophosphate (GMP), cytidine monophosphate (CMP) and uridine monophosphate (UMP) (**Fig.S5B**). Surprisingly, we observed an increase in FBP levels in *yap*-/- mutant embryos, however, this may be due to allosteric activation of phosphofructokinase (PFK) activity by elevated AMP (Bruser et al. 2012). Collectively, the decrease in glycolytic intermediates and concomitant increase in nucleotide monophosphates could be a manifestation of energetic stress, caused by abnormal glucose metabolism.

To precisely examine glucose uptake during development, the fluorescent glucose analogue GB2 was visualized in WT and *yap*-/- mutant larvae *in vivo* at 5 dpf, revealing suppressed GB2 uptake in *yap*-/- mutants (**Fig.S5C,D**). To gain greater insight into the fate of glucose *in vivo*, ^13^C relative isotopic enrichment was assessed after exposing WT and *yap*-/- mutant embryos to [U-^13^C_6_]-glucose from 48-72 hpf. Isotopic enrichment in the glycolytic intermediates G6P, DHAP and G3P was significantly reduced in *yap*-/- mutants at 72 hpf, supporting a lower rate of glucose uptake, without changes in isotopic enrichment of the downstream TCA cycle intermediates α-ketoglutarate and oxaloacetate (**Fig.5F; S5E**). Importantly, we observed a dramatic reduction in isotopic enrichment of intermediates in *de novo* biosynthesis of nucleotides in *yap*-/- mutants such as carbamoyl-L-aspartate, cytosine and guanosine (**Fig.5G**). Collectively, these results illustrate that Yap plays a critical role in the uptake and utilization of glucose for de novo nucleotide synthesis.

### Metabolic intervention impacts Yap-driven growth phenotypes

Given the dramatic decrease in nucleotide biosynthesis observed in *yap*-/- mutant embryos, we tested whether nucleoside replacement, achieved through aqueous exposure to a cocktail of deoxynucleosides (NS: deoxyadenosine (dA), deoxyguanosine (dG), deoxycytidine (dC), thymidine (dT)) could rescue aspects of the Yap-deficient phenotype. NS exposure of *yap*-/- embryos from 1 to 3 dpf significantly decreased the incidence of cardiac edema (observed in severely impacted fraction of *yap*-/- embryos, **Fig.S1A**) from 17.4% to 12.9% (**Fig.6A**) and increased liver size in *yap*-/- embryos, while demonstrating slightly toxic effects on liver growth in WT embryos (**Fig. S6A,B**). These results indicate a functional link between Yap, glucose transport and nucleotide biosynthesis for normal development and organ growth. To extend our findings beyond the examination of Yap loss on glucose metabolism and liver growth and test whether Yap-driven liver growth was dependent on *glut1*, we examined *lf:Yap* larvae, expressing an activated form of Yap in hepatocytes that causes increased liver growth by 5 dpf (Cox et al. 2016a). Analysis of liver size in WT and *lf:Yap* larvae following exposure to the irreversible Glut1 inhibitor WZB117 (herein referred to as WZB) (Liu et al. 2012; Shibuya et al. 2015) from 3-5 dpf demonstrated that hepatomegaly in *lf:Yap* larvae was effectively suppressed by WZB, whilst having no effect on WT larvae (**Fig.6B,C**). To corroborate these findings, we examined the effect of the competitive Glut inhibitor trehalose (DeBosch et al. 2016). Similarly to WZB, trehalose exposure significantly suppressed hepatomegaly in *lf:Yap* larvae, without affecting WT larvae (**Fig.6D**). These studies demonstrate that Yap-driven liver growth conditionally requires GLUT1-mediated uptake of glucose and its utilization for *de novo* nucleotide biosynthesis, and reveals potential therapeutic targets to inhibit Yap-mediated liver growth during carcinogenesis.

**Figure 6:**
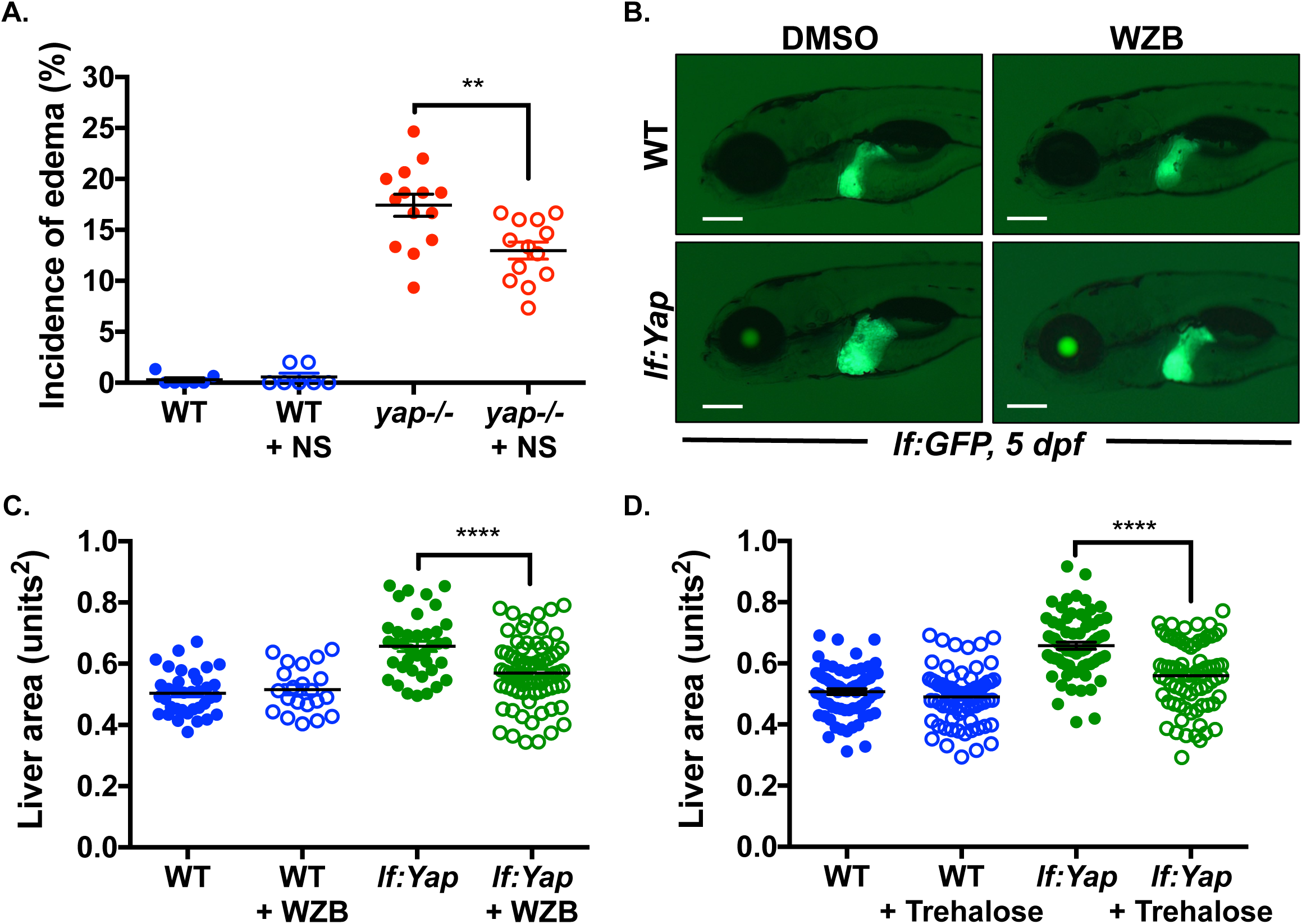
Metabolic intervention impacts Yap-driven growth phenotypes. A) Effect of nucleoside (NS) cocktail treatment from 1-3 dpf on the incidence of cardiac edema in WT and *yap*-/- mutant larvae. Each point represents the percentage of affected embryos in each independent clutch. n>7, ^∗∗^p<0.01, two-sided Student’s t-test; values represent the mean s.e.m. B) Effect of WZB-117 (WZB, 10 μM) treatment from 3-5 dpf in WT and *If:Yap* larvae on liver size as determined by fluorescence microscopy at 5 dpf in *lf:GFP* reporters. Scale bar: 200μm. C) Quantitative analysis of the effect of WZB treatment from 3-5 dpf on liver size as determined by fluorescence microscopy. n>21, ^∗∗∗∗^p<0.0001, two-sided Student’s t-test; values represent the mean ± s.e.m. D) Quantitative analysis of the effect of Trehalose (100 μM) treatment from 3-5 dpf on liver size as determined by fluorescence microscopy. n>70, ^∗∗∗∗^p<0.0001, two-sided Student’s t-test; values represent the mean ± s.e.m.

### The regulation of *glut1* expression and glucose uptake by Yap is evolutionarily conserved in mammals

To determine whether the regulation of glucose transporters and glucose uptake is conserved in mammals, gene expression data from the Cancer Cell Line Encyclopedia (CCLE) (Barretina et al. 2012) were analyzed, revealing a strong positive correlation between *GLUT1* (*SLC2A1*) mRNA and either *YAP1*, *AMOTL2* or *CYR61* expression (**Fig.7A,B; S7A**). In contrast, no correlation existed between *GLUT2* (*SLC2A2*) or *GLUT3* (*SLC2A3*) and either *YAP1*, *AMOTL2* or *CYR61* (**Fig.S7A**). Chromatin Immunoprecipitation sequencing (ChIPseq) datasets obtained from *LATS2* mutant MSTO211H cancer cells or HA-YAP^5SA^ expressing HuCCT1 liver cancer cells (Galli et al. 2015) revealed a strong TEAD-bound peak with YAP occupancy within an intragenic enhancer element of the *GLUT1* gene (**Fig.7C**). ChIP qPCR analysis verified enrichment of both YAP and TEAD at this enhancer element (**Fig.S7B**). The sequence of this enhancer element contains two consensus Tead Binding Site (TBS) motifs adjacent to a consensus AP-1 binding site (**Fig.S7C**), consistent with recent work demonstrating that YAP/TEAD co-occupy chromatin with AP-1 to drive expression of target genes (Zanconato et al. 2015). Furthermore, there was no evidence for binding of YAP or TEAD at the *GLUT2* or *GLUT3* locus (**Fig.S7D,E**). Together, these studies suggest that *Glut1* is directly regulated by Yap via an intragenic enhancer element.

**Figure 7:**
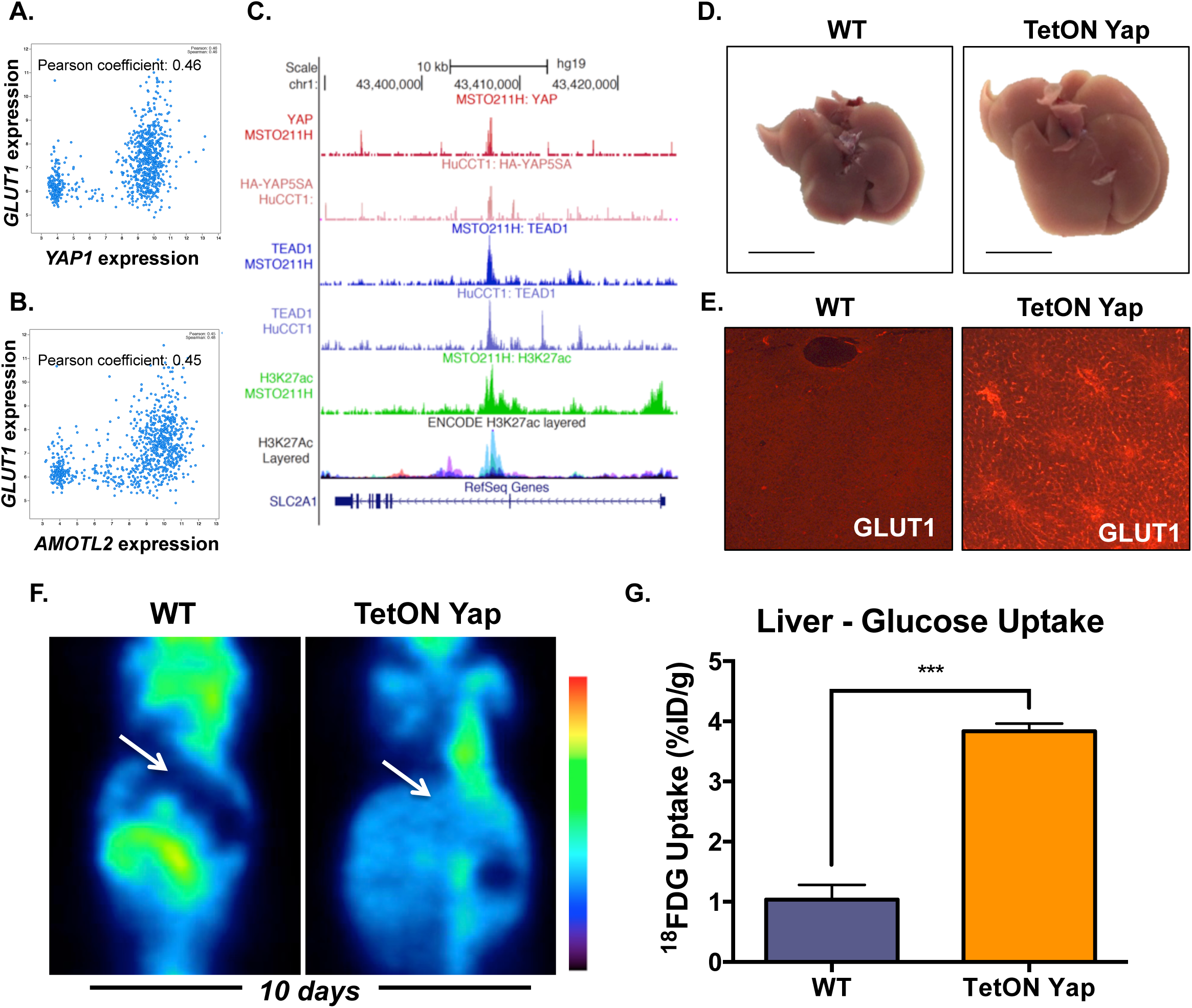
The regulation of *glut* expression and glucose uptake by Yap is evolutionarilv conserved. A) Dot plots showing the correlation between *GLUT1* and *YAP1* mRNA expression in the cancer cell line encyclopedia (CCLE, 967 lines). Pearson coefficient is 0.46. B) Dot plots showing the correlation between *GLUT1* and *AMOTL2* mRNA expression in the cancer cell line encyclopedia (CCLE, 967 lines). Pearson coefficient is 0.45. C) ChIPseq tracks at the *GLUT1* locus highlighting YAP and TEAD enrichment overlaying H3K27Ac marks at an intragenic enhancer element in different contexts (*LATS2* mutant MST0211H cancer cells and HA-YAP^5SA^ overexpressing HuCCTI liver cancer cells). D) Gross morphology of the liver from WT and TetONYap transgenic mice, treated with doxycycline for 10 days. Scale bar: 1 cm. E) Immunohistochemical analysis of GLUT1 expression in liver tissue from WT and TetONYap transgenic mice, treated with doxycycline for 10 days. Scale bar: 50μm. F) FDG PET imaging of coronal views from WT and TetONYap transgenic mice, treated with doxycycline for 10 days. Arrow is pointing to the liver. G) Scintillation analysis of FDG uptake (fraction of injected dose per gram of liver tissue, %ID/g) in dissected liver tissue from WT and TetON Yap transgenic mice, treated with doxycycline for 10 days. n=4, ^∗∗^p<0.01, two-sided Student’s t-test; values represent the mean ± s.e.m.

To determine whether the functional impact of Yap on glucose metabolism is evolutionarily conserved in mice, we used a previously described hepatocyte-specific Yap transgenic mouse model (TetONYap) (Yimlamai et al. 2014), enabling the conditional expression of activated Yap. Administration of doxycycline to TetONYap mice for 10 days leads to the onset of hepatomegaly (**Fig.7D**). Under these conditions, expression of the Yap-target gene *Ctgf* was elevated 12-fold, whilst expression of *Glut1* increased 3-fold (**Fig.S7F,G**). Corroborating the expression data, a significant increase in the amount of Glut1 at the plasma membrane of hepatocytes was detected by immunohistochemical analysis of TetONYap mouse livers (**Fig.7E**). To investigate a potential functional consequence on glucose uptake, we examined 18F-fluorodeoxyglucose (^18^FDG) uptake in WT and TetONYap mice at 10 days post doxycycline administration, prior to the onset of malignancy, was assessed by PET imaging and scintillation counting of dissected liver tissue. TetONYap liver tissue was FDG-avid, exhibiting a 4-fold increase in ^18^FDG uptake (**Fig.7F,G**). Collectively, these studies reveal that the regulation of glucose transporter expression and glucose uptake by Yap is evolutionarily conserved.

## Discussion

In the current study, we discovered defects in liver growth and glucose homeostasis in *yap*-/- mutant zebrafish that persist into adulthood. Transcriptional profiling revealed that Yap is both necessary and sufficient to regulate expression of glucose transporters *glut1* and *glut2.* Consistent with these changes, metabolic analyses demonstrated that adult *yap*-/- mutant zebrafish were glucose intolerant, with decreased glucose uptake and utilization in glycolysis and reduced *de novo* nucleotide biosynthesis. In addition, Glut1 activity was conditionally required for Yap-driven hepatomegaly. The direct regulation of glucose uptake by Yap is conserved in the murine liver, prior to the onset of malignancy. Collectively, our studies provide novel mechanistic insight into Yap-driven tissue growth that may have important therapeutic implications for targeting Yap-dependent growth in cancer.

### Role of Yap in stem cell homeostasis and organ development

Many fundamental insights into the role of Yap in organ development have been elucidated by murine studies. Previous work showed that *Yap*-/- mice are embryonic lethal at E8.5 (Morin-Kensicki et al. 2006). Subsequent studies demonstrated that conditional loss of *Yap* in differentiated liver tissue impaired liver function, by inhibiting bile duct formation, elevating hepatocyte cell death and promoting the development of steatosis and fibrosis (Zhang et al. 2010). Adult Yap-deficient livers are more susceptible to bile duct ligation-induced liver injury, exhibiting increased hepatocellular necrosis and decreased regeneration of cholangiocytes and hepatocytes (Bai et al. 2012). In the current study, we provide novel mechanistic insight into how Yap controls liver size by demonstrating that the observed defects in liver growth and hepatic progenitor potential *yap*-/- zebrafish are linked to impaired glucose uptake and subsequent use of glucose to support deoxynucleotide synthesis. Importantly, these defects can be partially rescued by repletion of deoxynucleotide pools. Loss of *Yap* has been shown to impede growth and dysregulate progenitor cell differentiation in a number of different tissues, including the small intestine (Cai et al. 2010; Barry et al. 2013), the heart (Xin et al. 2011; Xin et al. 2013), the kidney (Reginensi et al. 2013; Reginensi et al. 2015), the mammary gland (Chen et al. 2014) and the neural crest (Wang et al. 2016). It will be of great importance in future studies to determine whether perturbations in glucose and nucleotide metabolism contribute to the defects in progenitor cell fate observed in Yap-deficient organs other than the liver.

### Regulation of anabolic glucose metabolism in organ growth and cancer

Increased glucose uptake is a hallmark of tumor cells that enables the use of FDG-PET imaging to aid in the detection and prognosis of tumors as well as monitoring their response to therapy (Vander Heiden et al. 2009). Clinical studies reveal that FDG uptake in patients with hepatocellular carcinoma (HCC) is correlated with *GLUT1* expression (Mano et al. 2014). Although FDG-PET scans are routinely used to detect tumors, studies in transgenic mice have found that FDG uptake is not correlated with the rate of tumor growth or cellular proliferation; rather, FDG uptake is oncogene-specific, with Akt1 and Myc-driven tumors exhibiting higher FDG uptake than Wnt, Her2/neu or Ras-driven tumors (Alvarez et al. 2014). Consequently, it is essential to understand the molecular mechanisms by which tumor cells stimulate glucose uptake. In the current study, we have identified a previously unappreciated role for the Hippo pathway effector Yap in regulating glucose uptake by directly regulating the expression of *Glut1* via an intragenic enhancer element. Strikingly, the regulation of glucose uptake by Yap occurs prior to the onset of malignancy. It will be of interest in future studies to examine how the interplay between the Hippo pathway and other oncogenic pathways impact on glucose metabolism in the context of cancer.

### The Hippo/Yap pathway acts as a nutrient sensor coordinating metabolism and growth

Recent studies have shed light on the mechanisms by which the Hippo pathway effector Yap is regulated by nutrients in the cellular environment (Santinon et al. 2015). The first evidence for this came from the discovery that serum-borne lipids, such as lysophosphatidic acid and sphingosine 1-phosphate, regulate Yap activity (Miller et al. 2012; Yu et al. 2012). Further work expanded these findings by demonstrating that lipid products of mevalonate and bile metabolism can also promote Yap activation (Anakk et al. 2013; Sorrentino et al. 2014; Wang et al. 2014). Recently, a link between glucose concentration and Hippo pathway activity has emerged, wherein glucose deprivation or bioenergetic stress activates AMPK and LATS1/2, which phosphorylate and inactivate Yap (DeRan et al. 2014; Enzo et al. 2015; Gailite et al. 2015; Mo et al. 2015; Wang et al. 2015). The importance of lipid and glucose levels in the regulation of the Hippo pathway is best exemplified by elegant work demonstrating the activation of Tead and Yap directly by palmitoylation (Chan et al. 2016) and O-GlcNAcylation (Zhang et al. 2017), respectively. Together, these studies indicate that Yap activity is tightly regulated by the energetic state of the cell, particularly lipid and glucose supply.

Given the extent to which the Hippo pathway senses nutrient status, there is intense interest in identifying whether Yap reprograms metabolic output to enhance tissue growth. We have previously used metabolic and transcriptomic profiling to discover that Yap directly induces glutamine synthetase (*glul*) and reprograms glutamine metabolism towards *de novo* nucleotide biosynthesis to fuel liver growth and tumorigenesis (Cox et al. 2016a). In the current study, we use a combination of approaches to reveal that Yap directly induces the glucose transporters to redirect glucose utilization towards anabolism. The switch to anabolic glucose metabolism is supported by the recent demonstration that Yap is able to repress gluconeogenesis via suppression of PGC1α (Hu et al. 2017). In addition to our studies elucidating how Yap regulates glucose transporters, recent reports demonstrate that Yap directly induces the expression of the amino acid transporters SLC38A1 and SLC7A5 to stimulate cell growth (Hansen et al. 2015; Park et al. 2016). Together, these studies form the basis of an intriguing hypothesis that the Hippo pathway senses nutrients in the environment, whilst Yap mediates metabolic adaptation by co-ordinating biosynthetic output with tissue growth. Under conditions of nutrient deprivation, Yap is inhibited, which suppresses the energy consuming process of tissue growth. However, in nutrient rich conditions, Yap induces the expression of genes that stimulate nutrient uptake and biosynthesis, which facilitates tissue growth.

Although much of the attention on the Hippo pathway effector Yap has focused on its hyperactivation in the context of cancer, few studies have examined the concept that Yap deficiency leads to metabolic dysfunction. Our studies clearly demonstrate that the loss of Yap not only impacts glucose uptake, but more broadly disconnects nutrient sensing by the Hippo pathway, leading to disruption of metabolic homeostasis, as evident from the glucose intolerance in the Yap mutant fish. In line with this notion, Yap is known to regulate a network of genes involved in pancreas development (Cebola et al. 2015), defects in which could lead to metabolic dysfunction. Furthermore, elegant studies by Ardestani et al. have revealed that the Hippo kinase MST1 is a key regulator of β-cell dysfunction in diabetes (Ardestani et al. 2014). In this study, the authors showed that MST1 was activated in patients with diabetes and caused β-cell death, which could be rescued by loss of MST1. In light of these findings, we speculate that activation of the Hippo pathway and resulting decrease in Yap activity may play an important role in metabolic dysfunction and diabetes. Yap may influence metabolic homeostasis and disease via two broad mechanisms. First, Yap can impact the development of organs and tissues that have central roles in regulating systemic glucose homeostasis, such as the pancreas or liver. Second, Yap can regulate glucose uptake and metabolism in a cell-autonomous manner.

In summary, we have discovered an evolutionarily conserved role for Yap in the regulation of glucose homeostasis, regulating Glut transporter expression and glucose uptake. Metabolic profiling revealed that Yap regulated the anabolic utilization of glucose for *de novo* nucleotide biosynthesis. Importantly, we found that glucose uptake was conditionally required for Yap-driven liver growth.

Collectively, these studies provide insights into the mechanism by which Yap reprograms glucose metabolism and thereby identify a metabolic dependency of Yap-driven tissue growth on glucose transport that could be exploited to combat liver cancer.

## Materials and Methods

### Experimental procedures

#### Zebrafish husbandry

Zebrafish were maintained according to institutional animal care and use committee (IACUC-BIDMC) protocols. Lines used in this study include WT (AB), *yap*-/-^*mw48*^ mutant (Miesfeld et al. 2015), *taz*-/-^*mw49*^ mutant (Miesfeld et al. 2015), Tg(−*2.8fabp10a:yap1*^*S87A*^; –*0.8cryaa:Venus*)^*s705*^, abbreviated *lf:Yap* (Cox et al. 2016a), Tg(*fabp10a:CFP*-*NTR*)^*s931*^, abbreviated *lf:CFP* (Choi et al. 2014) and Tg(-*2.8fabp10:EGFP*)^as3^, abbreviated *lf:GFP* (Her et al. 2003). For this study, we generated Tg(*hsp70:mcherry*-*2A*-*flag*-*yap1*^*S87A*^), referred to as *hs:Yap* and Tg(*hsp70:dnyap*-*2A*-*mcherry*), referred to as *hs:dnYap* using standard Tol2 transgenesis techniques (Kwan et al. 2007). We used plasmids pME-YAPS87A and pME-NLS-YapDN generously provided by Dr. Brian Link (Medical College of Wisconsin, Milwaukee, WI) to generate the heat shock inducible transgenics. YAPS87A was amplified from pME-YAPS87A using the forward (5’-AAA AGA ATT CAg act aca aag acg atg acg aca agG ATC CGA ACC AGC ACA ACC-3’) and reverse (5’-AAA AAG AAT TCC TAT AGC CAG GTT AGA AAG TTC TCC-3’) primers to add a Flag-tag (lower case) and EcoRI restriction digest sites (underline). The resulting product was digested with EcoRI-HF (NEB) and cloned into the Tol2 kit vector p3E-2A-pCS2MCS-pA previously digested with EcoRI-HF and calf intestine phosphatase treated (Kwan et al. 2007). The resulting plasmid was recombined by Gateway (Life Technologies) recombination with Tol2 kit plasmids pDestTol2pA, p5E-hsp70i and pME-mCherry-nostop to place the heat shock cognate promoter (hsp70i) upstream of the constitutively active YAPS87A followed by a 2A self cleaving linker sequence and mCherry reporter gene. To create the Tg(hs:dnYAP-2A-mCherry), QuikChange lightning site-directed mutagenesis (Agilent Technologies) was used to delete the stop codon from the plasmid pME-NLS-YapDN with the oligos 5’-AGA AGG AGA GAC TGA GGA ACC CAG CTT TCT TGT A-3’ and 5’-TAC AAG AAA GCT GGG TTC CTC AGT CTC TCC TTC T-3’. The resulting plasmid pME-NLS-YapDN-nostop was recombined by Gateway cloning with Tol2 kit plasmids pDestTol2pA, p5E-hsp70i and p3E-2A-mCherrypA. To generate stable transgenic lines, we co-injected our newly generated plasmids with Tol2 Transposase RNA into one-cell AB embryos. Adult founders (F0) were identified by screening their progeny for heat shock inducible mCherry expression. The F1 generation’s progeny were then screened for Mendelian inheritance, inducible mCherry expression and brightness, and quantification of mCherry Hippo target gene mRNA expression.

#### Mouse Lines

Tetracycline-inducible *Yap^S127A^* expressing mice were previously described (Camargo et al. 2007; Yimlamai et al. 2014). AAV8-TBG-Cre (University of Pennsylvania Vector Core, AV-8-PV1091) was given to 4- to 8-week-old mice retro-orbitally. Male and female mice were used in this study and did not show gender-dimorphic differences. Transgene expression was induced by replacing normal drinking water with 5% sucrose containing doxycycline (1 mg/ml), whereas control (WT) siblings were given 5% sucrose water alone.

#### Genotyping

*yap*-/- and *taz*-/- mutants were genotyped as previously described (Miesfeld et al. 2015). Briefly, genomic DNA was isolated from zebrafish tissue using NaOH(Meeker et al. 2007). *yap* was amplified by PCR (forward primer: 5’-AGTCATGGATCCGAACCAGCACAA-3’, reverse primer: 5’-GCAGGCTGAAAGTGTGCATTGCC-3’) and the 4 bp deletion mutant was assessed for the presence of a Tfi1 restriction site. Similarly, *taz* was amplified by PCR (forward primer: 5’-CTCGGCTGAAACTACTTAAGGACG-3’, reverse primer: 5’-CTAAACAGTGTGCAGGAATGTCC-3’) and the 5 bp deletion mutant was assessed for the presence of a Hinf1 restriction site.

#### Heat shock conditions

Embryos were heat shocked by transferring them into pre-warmed (38 °C) egg water and incubating them at 38 °C for 30 min. Genotype was determined by the presence of mCherry fluorescence at 3 hours post heat-shock. Sorted non-fluorescent (WT) siblings were used as controls.

#### Fluorescence determination of liver size

*lf:CFP* of *lf:GFP* larvae were anesthetized with 0.04 mg/ml tricaine methane sulfonate (MS-222) and imaged by fluorescence microscopy using a Zeiss Discovery V8/Axio Cam MRC. Liver area was quantified using FIJI (NIH) as previously described (Schindelin et al. 2012; Weber et al. 2014).

### Histology and immunohistochemistry

Paraformaldehyde (PFA)-fixed larvae, fish were paraffin-embedded, serially sectioned and stained with Hematoxylin and Eosin (H&E), Periodic Acid-Schiff (PAS), Masson’s Trichrome or Reticulin stain using standard protocols (Goessling et al. 2008). Slides were deparaffinized and rehydrated prior to heat-induced antigen retrieval. Glucose transporter 1 (GLUT1) expression was detected using a 1:200 dilution of rabbit α-GLUT1 monoclonal antibody (Abcam, EPR3915) in conjunction with a 1:500 dilution of Alexa Fluor 647-conjugated donkey α-rabbit secondary antibody.

### Whole mount in situ hybridization

Zebrafish embryos were fixed in 4% paraformaldehyde (PFA) at the specified stages, and in situ hybridization was performed according to established protocols (http://zfin.org/ZFIN/Methods/ThisseProtocol.html). RNA probes for *prox1*, *hhex*, *foxa3*, *gc*, *ifabp* and *trypsin* were used. Embryos were imaged in glycerol with a Zeiss Discovery V8/Axio Cam MRC with the Axiovision software suite (Carl Zeiss). *prox1* expression areas were quantified using FIJI (NIH) as previously described (Schindelin et al. 2012; Weber et al. 2014).

### qRT-PCR

RNA was isolated from pooled zebrafish larvae or isolated livers using Trizol. Following DNAse treatment, cDNA was synthesized using the SuperScript III First-Strand Synthesis kit (Life Technologies). qRT-PCR was performed on biological triplicates using an iCycler with iQ SYBR green (BioRad). Gene expression was analyzed with *ef1a* as the reference gene (see Supplementary Table of primers).

### RNA transcriptomic analysis

RNA was extracted in Trizol (Life Technologies) WT and *yap*-/- mutant larvae at 3 dpf using the RNease Mini Kit (Qiagen), according to the manufacturers’ instructions. RNA quality was checked by Agilent Bioanalyzer Sequencing was performed after library construction on an Illumina HiSeq. polyA sequence data were annotated on the ZV9 genomic assembly to identify differentially affected genes (Collins et al. 2012), as previously described(Cox et al. 2016b). Gene ontology (GO) and Gene Set Enrichment analysis (GSEA) of biological processes was determined by GO Slim (Gene Ontology Consortium) and GAGE (Luo et al. 2009).

### Glucose tolerance test (GTT)

The GTT was performed in adult fish according to the methodology outlined by Eames et al(Eames et al. 2010). Zebrafish were fasted overnight, anaesthetized by brief hypothermic shock and given intraperitoneal injections of glucose (0.5 mg glucose per gram of fish weight) in Cortland salt solution (124.1 mM NaCl, 5.1 mM KCl, 2.9 mM Na_2_HPO_4_, 1.9 mM MgSO_4_ 7H_2_O, 1.4 mM CaCl_2_ 2H_2_O, 4% Polyvinylpyrrolidone and 10,000 U/L Heparin). Following specific periods of recovery, blood was collected following hypothermic shock by tail fin bleeding (Babaei et al. 2013). The concentration of glucose in collected blood was determined by an Accu-Chek Compact Plus (Roche Diagnostics) glucose meter according to the manufacturer’s instructions.

### Glucose uptake assays

Glucose uptake was examined by monitoring the uptake of the fluorescent glucose analogue GB2-Cy3, as previously described (Park et al. 2014). Zebrafish larvae were exposed to 20 μM GB2-Cy3 for 6 hr and imaged by fluorescence microscopy using a Zeiss Discovery V8/Axio Cam MRC. GB2-Cy3 uptake was quantified using FIJI (NIH) as previously described (Schindelin et al. 2012; Weber et al. 2014).

### Steady-state metabolomics analysis

Polar metabolites were isolated from WT and *yap*-/- mutant larvae at 3 dpf by methanol extraction. Polar metabolites were isolated and enriched using the methodology outlined by Yuan et al(Yuan et al. 2012). Metabolite fractions were collected and analyzed by targeted LC-MS/MS via selected reaction monitoring (SRM) with positive/negative ion polarity switching using a 5500 QTRAP hybrid triple quadrupole mass spectrometer (A/B SCIEX). Metabolic pathway enrichment analysis was performed using MetaboAnalyst (Xia et al. 2015).

### Analysis of ^13^C enrichment following [^13^C]-glucose exposure

WT and *yap*-/- mutant larvae were incubated with 1% [U^13^C_6_]-glucose at 2dpf for 24 hr. Larvae were subsequently washed and homogenized in methanol. Metabolites were quantified using SRM on a 5500 QTRAP mass spectrometer using a protocol to detect ^13^C-labelled isotopologues of polar metabolites, as previously described (Yuan et al. 2012).

### Chemical exposure

Zebrafish larvae were exposed to WZB-117 (WZB, 10 μM) from 3 to 5 dpf. Zebrafish embryos were exposed to a cocktail of nucleosides (NS, 25 μM deoxyadenosine, dA, 25 μM deoxyguanosine dG, 25 μM deoxycytidine, dC, 25 μM Thymidine, dT) from 1dpf to 3 dpf. Where necessary, chemicals were dissolved in DMSO. Chemicals were obtained from Sigma-Aldrich (USA).

### ChIP analysis

Chromatin Immunoprecipitation was performed as previously described (Galli et al. 2012; Galli et al. 2015). Cells were cross-linked in 1% formaldehyde for 10min at room temperature after which the reaction was stopped by addition of 0.125M glycine. Cells were lysed and harvested in ChIP buffer (100 mM Tris at pH 8.6, 0.3% SDS, 1.7% Triton X-100, and 5 mM EDTA) and the chromatin disrupted by sonication using a Diagenode Bioruptor sonicator UCD-200 to obtain fragments of average 200-500 bp in size. Suitable amounts of chromatin were incubated with specific antibodies overnight. Immunoprecipitated complexes were recovered on Protein-A/G agarose beads (Pierce) and, after extensive washes; DNA was recovered by reverse crosslinking and purification using QIAquick PCR purification kit (QIAGEN).

### PET and scintillation analysis of FDG uptake

Mice were administered doxycycline for 10 days and then fasted overnight prior to PET imaging. Mice received 200 μCi of FDG in 150 ml saline intravenously via the tail vein. Mice were subsequently anesthetized using isoflurane and PET images were acquired 1 hr post injection using a Siemens Focus 120 Scanner. PET images were reconstructed into a single frame using the 3D ordered subsets expectation maximization (OSEM3D/MAP) algorithm. After imaging, mice were euthanized by CO_2_ inhalation and liver tissue was collected, weighed and assessed for radioactivity by scintillation counting.

### Statistical analysis

All statistical analysis was performed using GraphPad Prism v7 (GraphPad Software, Inc.). Two-tailed unpaired Student’s t tests were performed for all experiments, unless otherwise specified. Results of independent experiments are presented as the mean ± SEM.

## Acknowledgements

This work was supported by an Irwin Arias Postdoctoral Fellowship (A.G.C.) and Liver Scholar Award (A.G.C.) from the American Liver Foundation, HDDC Pilot Feasibility Grant P30DK034854 (A.G.C.), NIH NIGMS T32GM007753 (K.L.H.), and by NIH NIDDK 1R01DK090311, 1R01DK105198, R24OD017870 (W.G.). W.G. is a Pew Scholar in the Biomedical Sciences and supported by the Claudia Adams Barr Program for Innovative Cancer Research. M.G.V.H. acknowledges support as an HHMI faculty scholar and additional support from SU2C and the NIH. E.C.L. is a fellow supported by the Damon Runyon Cancer Research Foundation. J.M.A. is partially supported by NIH NCI grants 5P01CA120964 and 5P30CA006516. G.G.G. is supported by an American-Italian Cancer Foundation postdoctoral research fellowship.

## Author Contributions

A.G.C. and W.G. designed experiments, reviewed results and wrote the manuscript. A.G.C. performed the majority of the zebrafish experiments assisted by A.T., K.L.H., M.F., and K.Y.M. J.M. and B.L. generated the *yap*-/- and *taz*-/- mutant fish. D.Y., G.G.G., B.H.F. and E.S. assisted with the murine studies. M.R.S., A.H., M.Y., K.K.B., E.C.L., M.L.S. and J.M.A. developed methods and assisted with metabolomics analysis. S.C. and Y.H. analyzed RNAseq datasets. J.A., Y.H., B.L., M.V.H., and F.D.C. provided overall input. All authors reviewed the manuscript.

---

**Table S1.**
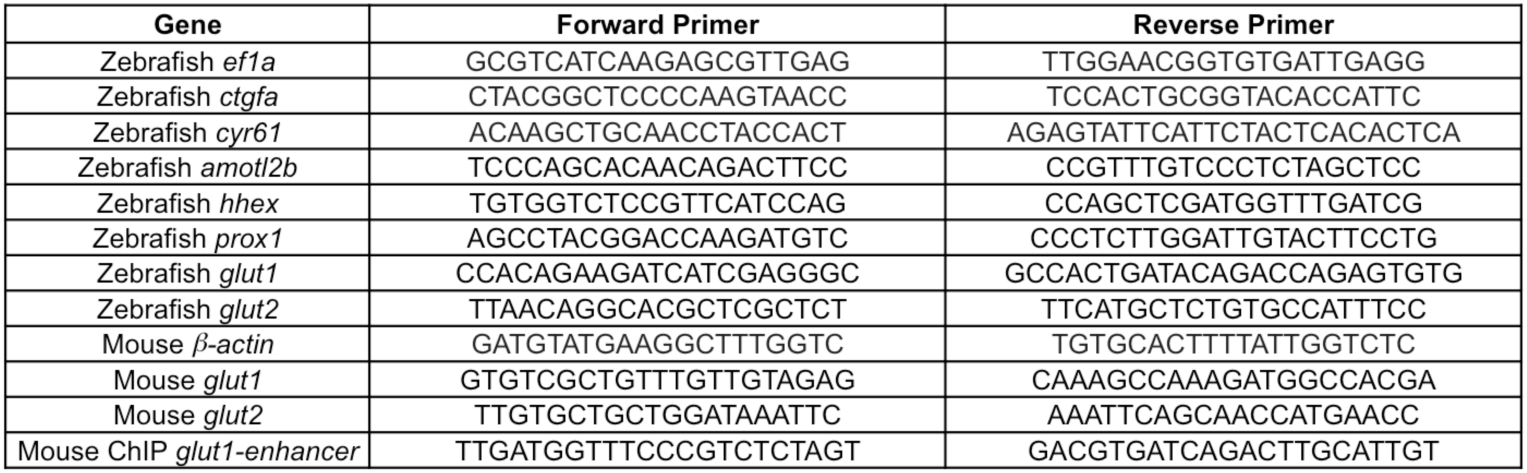
qRT-PCR Primers.

**Table S2.** Antibodies.

| Antibody | Supplier (order number) | Concentration |
| --- | --- | --- |
| α-Yap | CST (4912) | WB: 1/5000 |
|  | Clone gifted by Joseph Avruch, MGH, Boston | ChIP: 1/100 |
| α-H3K27ac | Abcam (ab4729) | ChIP: 1/1000 |
| α-IgG | Sigma (I8140) | ChIP: 1/100 |
| α-Tead4 | Abcam (ab58310) | ChIP: 1/100 |
| α-Glut1 | Abcam (epr3915) | IHC: 1/200 |
| α-Rabbit (Alexa Fluor®647) | Abcam (ab150075) | IHC: 1/500 |

